# Comprehensive Genomic Characterization of Breast Tumors with BRCA1 and BRCA2 Mutations

**DOI:** 10.1101/409813

**Authors:** Avantika Lal, Daniele Ramazzotti, Ziming Weng, Keli Liu, James M. Ford, Arend Sidow

## Abstract

**Background:** Germline mutations in the BRCA1 and BRCA2 genes predispose carriers to breast and ovarian cancer, and there remains a need to identify the specific genomic mechanisms by which cancer evolves in these patients. Here we present a systematic genomic analysis of breast tumors with BRCA1 and BRCA2 mutations.

**Methods:** We analyzed genomic data from breast tumors, with a focus on comparing tumors with BRCA1/BRCA2 gene mutations with common classes of sporadic breast tumors.

**Results:** We identify differences between BRCA-mutated and sporadic breast tumors in patterns of point mutation, DNA methylation and structural variation. We show that structural variation disproportionately affects tumor suppressor genes and identify specific driver gene candidates that are enriched for structural variation.

**Conclusions:** Compared to sporadic tumors, BRCA-mutated breast tumors show signals of reduced DNA methylation, more ancestral cell divisions, and elevated rates of structural variation that tend to disrupt highly expressed protein-coding genes and known tumor suppressors. Our analysis suggests that BRCA-mutated tumors are more aggressive than sporadic breast cancers because loss of the BRCA pathway causes multiple processes of mutagenesis and gene dysregulation.

## Background

Breast cancer is the most commonly diagnosed cancer and the second leading cause of cancer death among women. Approximately 10-15% of cases are associated with familial DNA repair-deficiency disorder, among which the most common forms are related to germline variants in BRCA1 and BRCA2, two genes involved in homologous recombination repair^1,2,3^. A germline mutation in BRCA1 or BRCA2 genes is known to be associated with a much higher than average lifetime risk (72% for BRCA1 and 69% for BRCA2 mutation carriers^4^) of developing breast cancer. In addition, these carriers also have a high risk of ovarian and other cancers^5,6^.

The reason for this is not entirely clear; however, two hypotheses are prevalent in the literature. As the BRCA1 and BRCA2 genes are both involved in the machinery for homologous DNA repair, it is hypothesized that defects in these genes leads to a lack of homologous repair activity, resulting in incorrectly repaired double-strand DNA breaks^7^. Incorrect repair of double-strand breaks can lead to higher rates of structural variation in the genome; the resulting structural variants may impact cell death or cell growth genes leading to cancer. In other words, an impaired BRCA complex could be a mutagen, analogous to environmental mutagens such as benzo(a)pyrene in tobacco smoke.

On the other hand, BRCA1/2-mutated tumors show a propensity to dedifferentiate into a more primitive state^8^, which could result in a higher rate of cell division. More cell division leads to accumulation of mutations as a result of DNA replication error, and these mutations may hit genes involved in cell death or growth leading to cancer. Under this hypothesis, lack of BRCA function is not a distinctive mutagen but an amplifier of normal mutational mechanisms^9^. Either of these phenotypes alone could explain the increased risk of cancer in BRCA mutation carriers but it is also possible for the two phenotypes to act synergistically. However, despite increasing literature on the topic, there has been no resolution, and the mechanisms underlying breast cancers in patients with BRCA mutations are still not fully comprehended.

Tumors with BRCA gene mutations often display a basal phenotype and are triple-negative (lacking ER, PR and HER2 amplifications)^10^. Previous studies have identified differences in point mutational signatures^11^, copy number profile^12^, gene expression signatures^13^ and patterns of structural variation^11^ between BRCA-mutated and sporadic breast tumors, indicating that tumor evolution follows a distinct path in these cancers. Moreover, in addition to patients with inherited germline mutations in the BRCA genes, somatic inactivation of BRCA1 and BRCA2 has also been reported in breast and ovarian tumors^14,15^. Recent studies suggest that such tumors may present similar phenotypes to those with germline BRCA inactivation^16^.

In this work we combine newly generated sequencing data with previous datasets, and perform an in-depth integrative analysis of genomic and epigenomic data in order to achieve better insights into the mechanism underlying tumor formation in individuals with BRCA gene mutations. Our aim here is to characterize the genomic variation in BRCA-mutated tumors and understand whether and how they are different from common classes of sporadic breast tumors. We present novel results on the differences in point mutation, DNA methylation, and structural variation in BRCA1/2 mutated tumors, and identify specific genes including known tumor suppressors that are frequently damaged by structural variation in these tumors.

## Results

### BRCA1/2-mutated tumors have a high burden of point mutations

To compare the point mutation profiles of BRCA1/2-mutated tumors with other breast tumors, we analyzed a published dataset of 560 breast tumors^11^. This dataset includes 36 tumors with inactivating mutations in BRCA1 (31 germline and 5 somatic mutations) and 39 tumors with inactivating mutations in BRCA2 (29 germline and 10 somatic mutations), as well as 118 triple-negative (TN) tumors, 293 ER+ (HER2−) tumors and 71 HER2+ tumors.

Both BRCA1-mutated and BRCA2-mutated tumors present significantly more genome-wide point mutations than the sporadic tumors (Wilcoxon test, p = 4.9×10^−11^ for BRCA1-mutated vs. sporadic tumors, p = 5.5×10^−11^ for BRCA2-mutated vs. sporadic tumors), with BRCA1-mutated tumors having a particularly high number of point mutations (Figure 1a); this phenomenon has been previously reported with smaller sample sizes^17^. Among the sporadic tumors, triple-negative (TN) tumors have higher mutation counts, close to those of BRCA2-mutated tumors – nevertheless, both BRCA1-mutated and BRCA2-mutated tumors have significantly higher mutation counts than the triple-negative tumors in this dataset (Wilcoxon test, p = 0.005 for BRCA1-mutated vs. TN, p = 0.019 for BRCA2-mutated vs. TN). Tumors with somatic BRCA1/2 mutations present similarly high mutation counts to those with germline inactivation.

**Figure 1.**
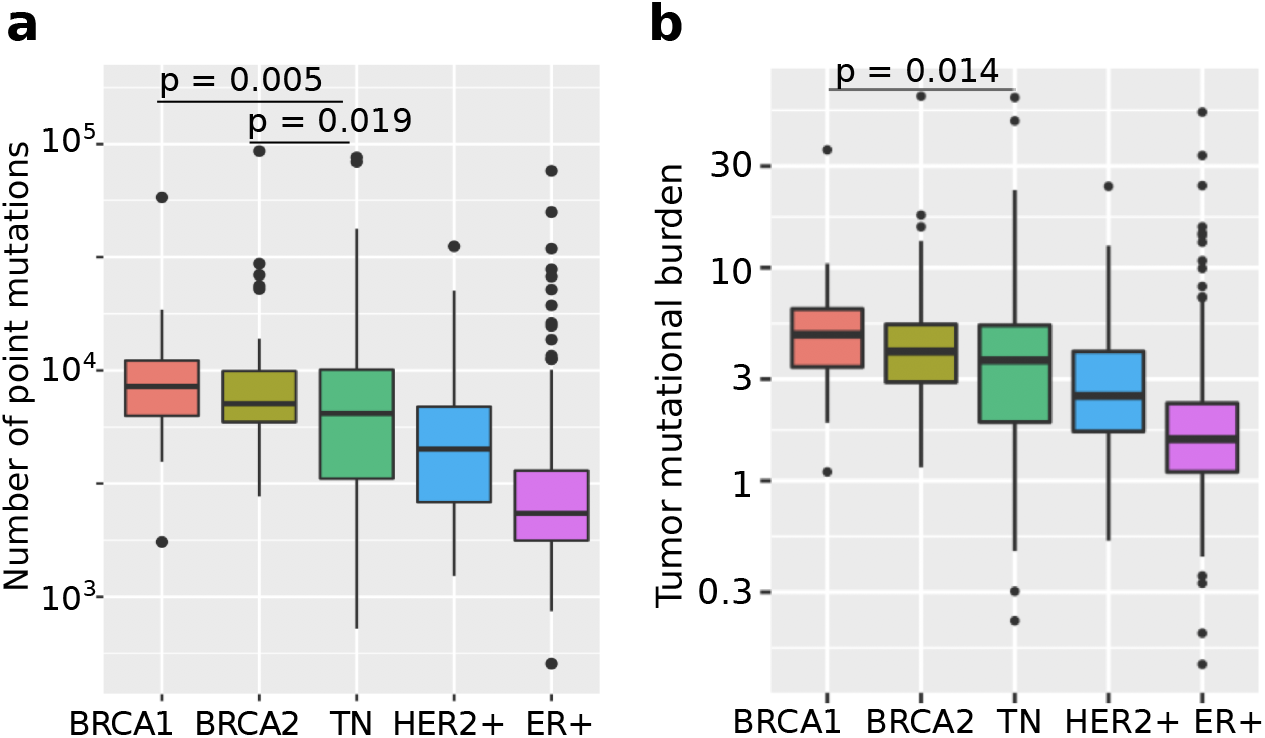
a) Boxplots showing the number of single nucleotide variants in the whole genomes of different classes of breast tumors, based on data from 560 breast tumors^11^. b) Boxplots showing the Tumor Mutational Burden (number of nonsynonymous mutations per Mb of coding sequence) in the whole genomes of different classes of breast tumors, based on data from 560 breast tumors^11^. TN = Triple-negative.

We also tested whether BRCA-mutated tumors have more missense mutations. Compared to sporadic tumors, both BRCA1-mutated and BRCA2-mutated tumors show a higher Tumor Mutational Burden (TMB), calculated as the number of nonsynonymous mutations per Mb of coding sequence (Figure 1b; Wilcoxon test, p = 9.9×10^−10^ for BRCA1-mutated vs. sporadic tumors, p = 5.1×10^−8^ for BRCA2-mutated vs. sporadic tumors). However, in this case, the triple-negative tumors in our dataset are not significantly different from the BRCA2-mutated tumors.

### Differences in mutational signature exposures between BRCA1/2-mutated and sporadic tumors

Mutational signatures are patterns of point mutations in the genome created by specific mutagenic processes, e.g., a chemical mutagen or a defect in a DNA repair enzyme^18^. If BRCA1/2-mutated tumors evolve via distinct point mutation-causing processes, they may possess unusual mutational signatures. We therefore analyzed whether the BRCA1/2-mutated tumors have a different pattern of mutational signatures from the sporadic breast tumors.

A previous study^11^ applied a widely used framework^18,19^ for extracting mutational signatures from genomic data to the same dataset of 560 breast tumors, resulting in 12 mutational signatures. Notably, the resulting signatures are very dense, and many are also very similar to each other. While some have been linked to known mutational processes in breast cancers, others still have no known etiology^20^. This may be due to the fact that this framework extracts as many signatures as required to improve the fit to the data, without testing whether these signatures perform well at fitting unseen data. This can be expected to result in a high number of signatures that potentially overfit the data. For these reasons, we wished to use a principled approach that incorporates biological knowledge, as well as statistical methods to prevent overfitting.

We recently developed SparseSignatures^21^, a novel framework to identify mutational signatures. This method incorporates a background model representing the pattern of mutations caused in the normal course of cell division by DNA replication errors - a signature that we assume is present in all tumors. The background signature is fixed and additional signatures are discovered while incorporating a LASSO constraint to ensure that the signatures are sparse, producing a more biologically accurate and interpretable solution. SparseSignatures also applies a repeated bi-cross-validation strategy^21^ to select the number of signatures. This allows us to avoid overfitting by selecting a number of signatures that not only fit the data used to discover them but are also capable of predicting unseen data points.

We applied this approach to 555 breast tumors (we removed 5 tumors with <1000 mutations as previously described^21^). We discovered 8 mutational signatures in addition to the background (Figure 2a, Supplementary Table 1). These signatures are statistically strongly supported and most of them are related to known mutagenic mechanisms. We compared our discovered signatures to previously proposed signatures in the COSMIC database based on cosine similarity and visual inspection, and found that some of our signatures are highly similar to known signatures in COSMIC (Supplementary Tables 2 and 3.) Signature 1 is the well-known signature caused by deamination of methylated cytosines at CpG sites into thymine, and is highly similar to COSMIC signature 1. Signatures 2 and 5 are associated with deregulation of APOBEC cytidine deaminases^20^ and correspond to COSMIC signatures 2 and 13 respectively. Signature 3is similar to COSMIC signature 12 previously associated with defective DNA mismatch repair^22^. Signature 4 is a pattern of elevated TT>GT point mutations, highest in a CTT context, and corresponds to COSMIC signature 17.

**Figure 2.**
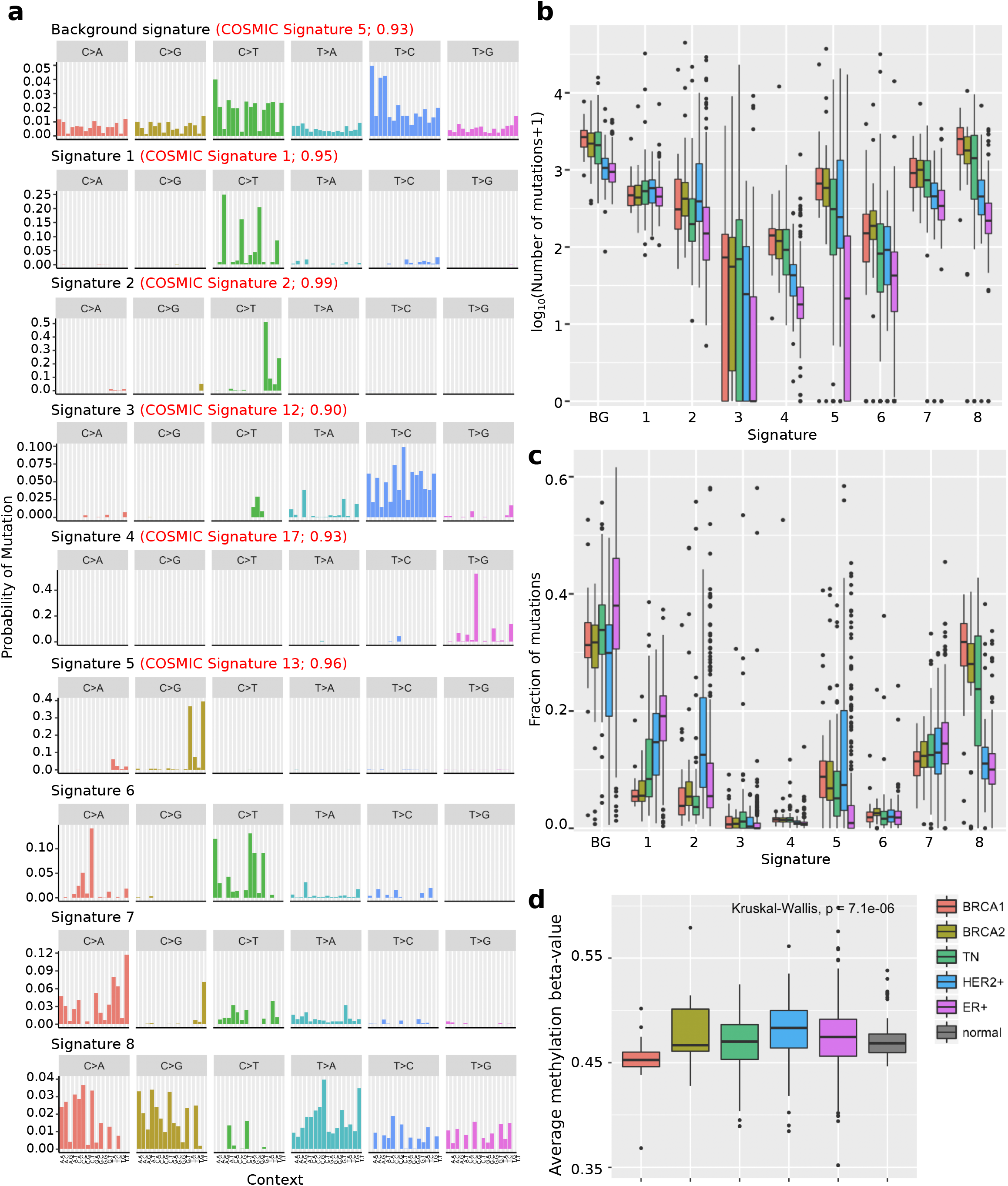
a) 9 signatures (including background) discovered by applying SparseSignatures to the whole genomes of 555 breast tumors. For signatures 1-5, the corresponding COSMIC signature is listed in red along with its cosine similarity to the discovered signature. b) Boxplots showing the number of mutations attributed to each signature in different classes of breast tumors. c) Boxplots showing the fraction of mutations attributed to each signature in each sample, for different classes of breast tumors. d) Average beta-value (representing the extent of cytosine methylation) of CpG sites, in different classes of breast tumors, based on data from TCGA^22^.

The remaining 3 signatures do not have high similarity to any of the signatures in COSMIC. Signature 6 may be associated with defective DNA mismatch repair based on common features with COSMIC signature 20, while Signature 4 is moderately similar to COSMIC Signature 18, which has recently been associated with DNA damage caused by reactive oxygen species^19^. Finally, Signature 8 is a relatively dense pattern characterized by an elevated rate of C>A, C>G and T>A mutations, whose etiology is unknown.

SparseSignatures also calculates the exposure values for each signature, i.e. the number of mutations originating from each signature in each patient (Supplementary Table 4). On average, the background signature, representing DNA replication errors, contributes more mutations than any other signature - an average of 1619.5 mutations per tumor, compared to the next highest value of 1017.6 mutations per tumor for signature 8. (Wilcoxon test, p = 1.27 × 10^−46^, exposure to background vs. exposure to signature 8). The higher number of point mutations in the BRCA1/2-mutated tumors, compared to sporadic tumors, is reflected in a higher exposure to the background signature (Figure 2b; Wilcoxon test, p=2.1 × 10^−17^). This suggests that BRCA1/2-mutated tumors have undergone more cell divisions.

We do not find any signature present only in BRCA1/2-mutated tumors. However, compared to sporadic tumors, BRCA1/2 mutated tumors have more mutations (Figure 2b; Wilcoxon test p=2.1 × 10^−20^) and a higher fraction of mutations (Figure 2c; Wilcoxon test, p=5.4 × 10^−18^) attributed to Signature 8. While the etiology of this signature is uncertain, it is not simply indicative of BRCA mutation as many sporadic triple-negative tumors also have a similarly high contribution by signature 8. In general, the mutational signature profiles of sporadic triple-negative tumors are very close to those of BRCA1/2-mutated tumors. This suggests that these tumors have similar underlying mutagenic processes, including a high contribution of the background signature indicating a high rate of cell division.

### BRCA1/2-mutated tumors have lower levels of CpG methylation

Signature 1 is underrepresented in BRCA1/2-mutated tumors; it contributes an average of 6.0% of the total mutations in BRCA1-mutated tumors and 6.6% in BRCA2-mutated tumors, as opposed to 16.0% of mutations in sporadic tumors (Figure 2c; Wilcoxon test, p = 7.6 × 10^−14^ for BRCA1 vs. sporadic, p = 2.1 × 10^−13^ for BRCA2 vs. sporadic). This signature is caused by DNA CpG methylation and subsequent deamination of methylated cytosine to thymine leading to C>T mutation. The ratio of Signature 1 exposure to background signature exposure is significantly lower in both BRCA1 and BRCA2-mutated tumors compared to sporadic tumors (Wilcoxon test, p = 9.2 × 10^−15^ for BRCA1 vs. sporadic and 8.6 × 10^−9^ for BRCA2 vs. sporadic; Supplementary Figure 1). Taking the background signature exposure as an indicator of cell division, this suggests that BRCA1/2-mutated tumors may have lower CpG methylation than sporadic tumors.

As DNA methylation data is not available for this dataset, we tested whether DNA methylation is lower in BRCA1/2-mutated tumors in a cohort of 674 breast cancers and 97 normal breast tissue samples from The Cancer Genome Atlas^23^. This dataset included 20 tumors with inactivating germline or somatic mutations in BRCA1 and 13 with inactivating germline or somatic mutations in BRCA2, as well as 114 HER2+ tumors, 417 ER+ (HER2−) tumors, and 110 triple-negative tumors (Supplementary Table 5). We found that global CpG methylation levels are indeed significantly reduced in BRCA1-mutated tumors compared to sporadic tumors (Figure 2d; Wilcoxon test, p = 3.1 × 10^−4^) and compared to normal tissue samples (Wilcoxon test, p = 3.3 × 10^−5^). On the other hand, there was no significant difference between BRCA1-mutated and sporadic tumors in the methylation level of the 3081 CpA sites measured on the same platform (Supplementary Figure 2).

We did not observe a significant difference in methylation levels between BRCA2-mutated and sporadic tumors. However, we note the low number of BRCA2-mutated samples in this analysis, which may limit our power. In order to test this, we performed a power analysis given the considered sample sizes. We first estimated the parameters of normal distributions that best fit the different groups from our data and then performed 10,000 simulations by drawing samples from these distributions. For each simulation, we tested whether we could correctly assess that the data were drawn from different distributions and estimated the power of the test as the fraction of simulations in which the results were correct. This analysis resulted in an estimated power of 0.27 when comparing BRCA2 tumours to normal tissues and of 0.16 when comparing BRCA2 tumours to sporadic ones.

### BRCA1-mutated tumors have elevated tandem duplications and interchromosomal translocations

We obtained whole-genome sequencing data for 67 of the 560 tumor samples^11^ along with their matched normal samples, for which BAM files were available for download from ICGC. In addition, we sequenced whole genomes from 14 additional breast tumors and matched normal samples^24^ from patients carrying germline BRCA1/2 mutations, resulting in a dataset of 81 tumor genomes: 27 with germline BRCA1 mutations, 19 with BRCA2 mutations (17 germline and 2 somatic), and 35 sporadic breast tumors without BRCA inactivation, of which 19 were triple-negative and 16 were ER+. Supplementary Table 6 describes the selected samples.

We used SvABA^25^ to identify somatic indels and structural variants in these tumor genomes. SvABA is a newly developed indel and structural variant caller that uses genome-wide local assembly and has been shown to possess superior sensitivity and specificity to previous methods^25^. After filtering the variant calls (see Methods), we identified a total of 7,234 high-confidence somatic indels and 19,684 high-confidence somatic structural variants in the 81 tumor genomes. We then compared BRCA1/2-mutated tumors against sporadic tumors. We included the 2 tumors with somatic BRCA2 inactivation along with those showing germline BRCA2 inactivation. We found that both BRCA1 and BRCA2-mutated tumors had significantly more indels (Wilcoxon test, p = 1.6 × 10^−5^ for BRCA1 and p = 1.4 × 10^−3^ for BRCA2) and structural variants (Wilcoxon test, p = 5.1 × 10^−7^ for BRCA1 and p = 0.029 for BRCA2) per tumor than the sporadic tumors.

We next examined specific types of variation. Both BRCA1 and BRCA2-mutated tumors have more deletions than sporadic tumors (Figure 3a; Wilcoxon test, p = 1.4 × 10^−6^ for BRCA1 and p = 1.5 × 10^−4^ for BRCA2). While most deletions in sporadic tumors are either <5 bp or >10 kb long, BRCA1/2-mutated tumors have a large number of deletions of intermediate size; the size distribution of these deletions is bimodal, with one peak between 5-100 bp and the other between 100 bp-10 kb (Figure 3b). While the 5-100 bp long deletions mostly lack microhomology at the breakpoints, the majority of the BRCA1/2-mutated samples have short regions of microhomology (1-10 bp) at the breakpoints in >50% of the deletions in the 100 bp-10 kb size range (Figure 3c).

**Figure 3.**
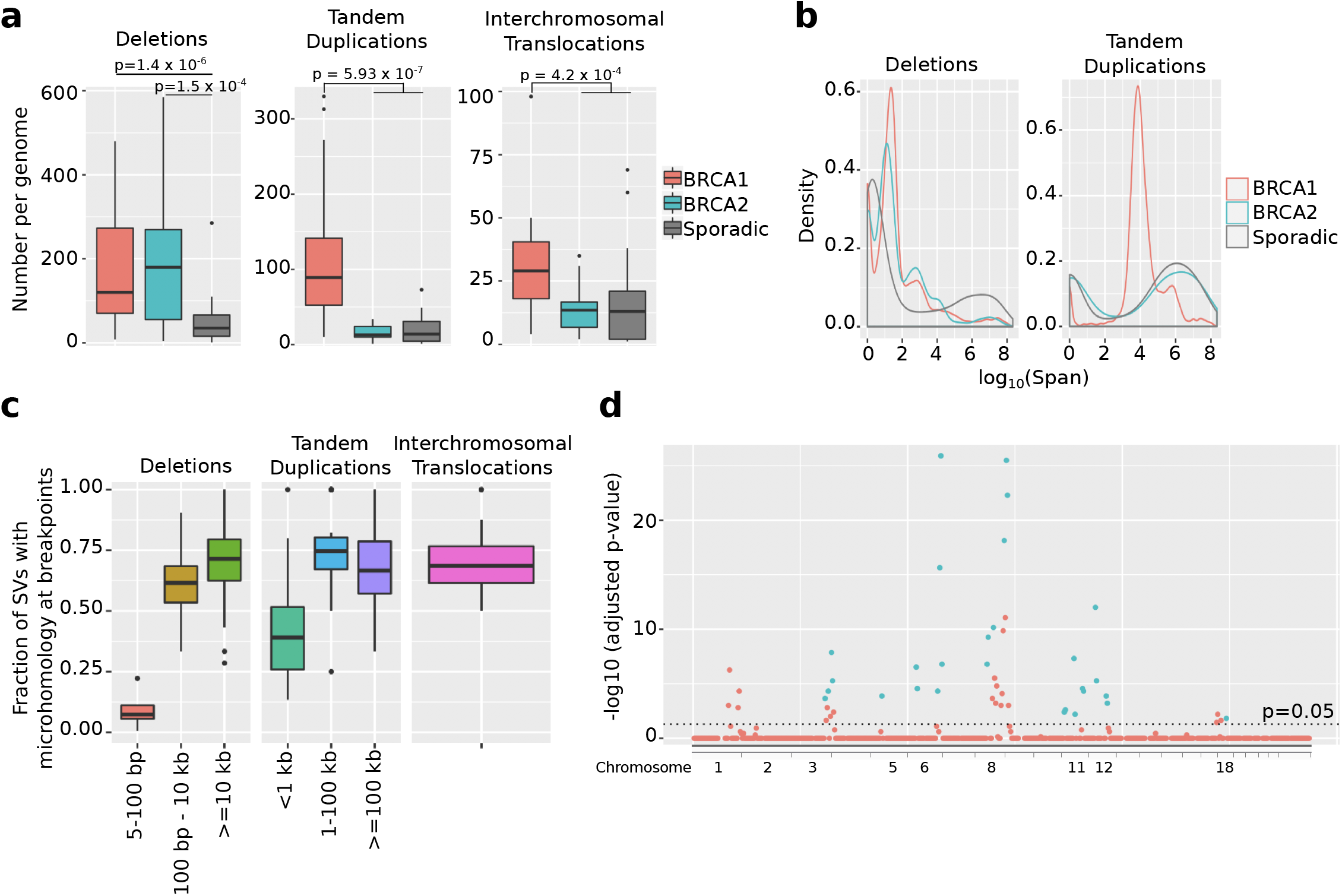
a) Boxplots showing the number of deletions, tandem duplications, and interchromosomal translocations, in the genomes of 81 breast tumors. b) Probability distributions of the sizes of deletions and tandem duplications in the genomes of 81 breast tumors. c) Boxplots showing the fraction of structural variants in a tumor genome that contain regions of microhomology at the breakpoint, divided into deletions, tandem duplications and interchromosomal translocations, for 46 BRCA1/2-mutated tumors. The x-axis shows size of the structural variants. d) Manhattan plot with the y-axis showing the bonferroni-corrected p-value for enrichment of structural variants in 10-Mb long genomic bins, for 46 BRCA1/2-mutated tumors. The y-axis shows the position of the bin. Chromosomes are ordered from 1 to 22 followed by X.

On the other hand, we observed that BRCA1-mutated tumors have an elevated number of tandem duplications^26^ (Figure 3a; Wilcoxon test, p(BRCA1 vs. others) = 5.9 × 10^−7^), predominantly ranging in size from 1-100 kb (Figure 3b). Most of the tandem duplications in this size range have short regions of microhomology at the breakpoints (Figure 3c).

In addition, we observed that the BRCA1-mutated tumors have more interchromosomal translocations than the BRCA2-mutated or sporadic tumors (Figure 3a; Wilcoxon test, p = 4.2 × 10^−4^). To our knowledge this phenomenon has not been described previously. Like the tandem duplications described above, these translocations also tend to have microhomology of 1-10 bp at the breakpoints (Figure 3c).

Large copy number alterations in the genome can significantly change genome size. To test whether the elevated numbers of point mutations and structural variants in BRCA1/2-mutated samples are due to biological differences or are accounted for by the availability of more DNA, we identified copy number variants in the genomes of these 81 tumor samples using Control-FREEC^27^. After correcting the size of the genome in each tumor to account for copy number alterations, we find that the BRCA1-mutated samples have larger genomes than BRCA2-mutated or sporadic tumors (Supplementary Figure 3, Wilcoxon test p = 3.2 × 10^−3^). However, normalizing the number of mutations for the actual size of the genome does not affect our results. BRCA1 and BRCA2-mutated tumors still have significantly higher numbers of point mutations and deletions than sporadic tumors, and BRCA1-mutated tumors have significantly higher numbers of tandem duplications and interchromosomal translocations than all other classes of tumors. BRCA2-mutated tumors are not significantly different from sporadic tumors in the number of tandem duplications or interchromosomal translocations.

### Functional regions hit by breakpoints in BRCA-mutated tumors

Since BRCA1/2-mutated tumors have an elevated number of structural variants, we tested whether these structural variants tend to disrupt functional and regulatory regions of the genome. We found that in BRCA1/2-mutated tumors, the breakpoints for interchromosomal translocations, 1-100 kb duplications and 100 bp-10 kb deletions are all enriched in regions of open chromatin (Hypergeometric test, p = 3.9 × 10^−6^ for interchromosomal translocations, p = 6.6 × 10^−6^ for 1-100 kb duplications and p = 0.034 for 100 bp-10 kb deletions).

Since BRCA1-mutated tumors have elevated numbers of tandem duplications and interchromosomal translocations, we examined these breakpoints specifically. The breakpoints for both 1-100 kb tandem duplications and interchromosomal translocations in BRCA1-mutated tumors are enriched in protein-coding genes (Hypergeometric test, p = 1.7 × 10^−14^ and p = 0.014 respectively). The breakpoints for 1-100kb tandem duplication breakpoints in BRCA1 tumors are also specifically enriched in exons (Hypergeometric test, p = 1.2 × 10^−3^). We also found that interchromosomal translocation breakpoints in BRCA1 tumors are enriched in TAD boundaries (Hypergeometric test, p = 2.9 × 10^−4^). Disruption of TAD boundaries has previously been shown to alter gene expression in tumors by modifying 3D contact domains on the chromosome^28^.

We also tested whether the indels and structural variant breakpoints in BRCA1/2-mutated tumors are associated with the local replication timing. The breakpoints for 5-100 bp long deletions and small (<5 bp) indels are both enriched in late replicating regions (Hypergeometric test, p = 4.3 × 10^−4^ and p = 9.6 × 10^−16^ respectively). On the other hand, the breakpoints for 1-100 kb tandem duplications and interchromosomal translocations, both of which are elevated in BRCA1-mutated tumors, are enriched in early replicating regions (Hypergeometric test, p = 4.6 × 10^−19^ and p = 3.2 × 10^−11^ respectively).

### Structural variants disrupt tumor suppressor genes

We examined the genes that are disrupted by indel and structural variation breakpoints in BRCA1/2-mutated tumors. The genes disrupted by both indels and SVs have significantly (Wilcoxon test, p < 10^−15^ for both) higher levels of expression in normal breast tissue, according to RNA-Seq data from GTEx^29^ (Supplementary Figure 4). Further, the set of genes disrupted by indels and structural variation are both significantly enriched for tumor suppressor genes (Hypergeometric test, p = 1.4 × 10^−5^ and p = 4.9 × 10^−10^ respectively).

We next searched for specific genes enriched for indels or structural variant breakpoints in the BRCA1/2-mutated tumors, using a poisson test. The null model here is that breakpoints are randomly distributed throughout the genome, and we identify protein-coding genes that have significantly more breakpoints than expected from their length. We identified 11 genes enriched for indels/structural variant breakpoints: NME7, KLHL8, EFNA5, PTEN, DHX32, ETV6, RB1, ARGLU1, TP53, P4HB, and RUNX1 (Table 1). After correcting the length of each gene to take into account its copy number in each tumor, 10 of these genes (KLHL8, EFNA5, PTEN, DHX32, ETV6, RB1, ARGLU1, TP53, P4HB, RUNX1) remained significant. Moreover, 4 of these genes (RB1, PTEN, KLHL8, and EFNA5) are also spanned by long deletions in multiple BRCA1/2-mutated samples, representing another mode of inactivation.

**Table 1:** 11 protein-coding genes show enrichment for indels/structural variant breakpoints in BRCA1/2-mutated tumors.

|  | Gene | Chromosome | Length (bp) | Number of indels/SV breakpoints | Adjusted p-value | Adjusted p-value (corrected for copy number) | Adjusted p-value (corrected for copy number) BRCA1 only | Adjusted p-value (corrected for copy number) BRCA2 only |
| --- | --- | --- | --- | --- | --- | --- | --- | --- |
| 1 | RB1 | 13 | 176159 | 17 | 1.31x10<sup>-9</sup> | **1.69x10<sup>-10</sup>** | 4.60x10<sup>-9</sup> | 1.00 |
| 2 | TP53 | 17 | 6986 | 5 | 5.48x10<sup>-5</sup> | **2.13x10<sup>-5</sup>** | 6.17x10<sup>-4</sup> | 1.00 |
| 3 | PTEN | 10 | 101523 | 10 | 1.45x10<sup>-4</sup> | **7.53x10<sup>-5</sup>** | 1.66x10<sup>-2</sup> | 1.00 |
| 4 | ETV6 | 12 | 240919 | 13 | 0.00102 | **0.00243** | 4.17x10<sup>-5</sup> | 1.00 |
| 5 | RUNX1 | 21 | 256765 | 13 | 0.00210 | **0.00275** | 4.73x10<sup>-5</sup> | 1.00 |
| 6 | KLHL8 | 4 | 32021 | 6 | 0.00372 | **0.00293** | $3.73 \times 10^{-4}$ | 1.00 |
| 7 | P4HB | 17 | 16460 | 5 | 0.00380 | **0.00592** | $5.62 \times 10^{-2}$ | 1.00 |
| 8 | NME7 | 1 | 234926 | 12 | 0.00581 | 0.0695 | $1.83 \times 10^{-3}$ | 1.00 |
| 9 | EFNA5 | 5 | 289359 | 13 | 0.0080 | **0.00193** | $3.28 \times 10^{-5}$ | 1.00 |
| 10 | ARGLU<br>1 | 13 | 23924 | 5 | 0.0233 | **0.0234** | $4.17 \times 10^{-3}$ | 1.00 |
| 11 | DHX32 | 10 | 44138 | 6 | 0.0236 | **0.0168** | 1.00 | 1.00 |

Aside from enrichment of breakpoints in genes and exons, we also wanted to test whether there are larger regions of the genome, including non-coding regions, that are enriched for indel or SV breakpoints in our set of 46 BRCA1/2-mutated tumor samples. These would include breakpoints for variants that span across whole genes, as well as those that affect gene expression by disrupting regulatory regions of the genome.

We divided the genome into 10-Mb long bins, overlapping by 5 Mb. We then combined all the high-confidence indels and structural variants collected from all the BRCA1/2-mutated tumors. We tested whether these tumors are enriched for indel/structural variant breakpoints in each bin using a poisson test, with the null model being that breakpoints are distributed uniformly across the genome. We found 48 bins that had a Bonferroni-corrected p-value of less than 0.05 (Figure 3d). All of these regions were disrupted by at least one indel or structural variant in at least 50% of BRCA1/2-mutated tumors. After correcting the number of bases in each bin to account for copy number changes, 28 bins remained significantly enriched (Bonferroni-corrected p<0.05). These bins are located on chromosomes 3, 5, 6, 8, 10, 11, 12, and 18, and several of them overlap with each other. Their coordinates are listed in Supplementary Table 7.

### Identification of large-scale structural variants on the basis of 10X Genomics data

Our analysis above, as well as previous studies^11,26^, highlight the importance of structural variation in the evolution of BRCA-mutated cancers. However, short-read sequencing is not ideal for accurate detection of large structural variants due to the limited read length. 10X Genomics is a linked-read technology, which uses barcodes to identify short fragments that originate from the same large molecules. Thus, it provides long-range information that offers improved resolution and detection of structural variants^30^. We sequenced additional DNA prepared with 10X technology from 3 tumors with BRCA1 germline mutations. In addition, we sequenced genomic DNA from 1 BRCA2-mutated tumor and 12 sporadic triple-negative tumors from the same study^23^. We used GROC-SVs^30^ to identify structural variants in these genomes and were able to confirm several translocations in the 3 BRCA1-mutated samples. Further, although the sample size is too small for a statistical test, we observed that these BRCA1-mutated samples had more translocations on average than the sporadic tumors (Supplementary Table 8).

Although structural variants are normally classified into simple categories (such as duplications, deletions, and translocations), recent studies have revealed that some tumor genomes also contain a large number of complex structural variants (CSVs) that cannot be explained by a simple end-joining or recombination event^31^. In our short-read data, we observe that 16% of structural variants are accompanied by a short insertion event at the breakpoint; the occurrence of such insertions is not significantly different in BRCA1/2-mutated tumors. However, larger CSVs composed of multiple rearrangements cannot be detected by short reads. Using GROC-SVs on the 10X data, we detected two complex structural variants in the sample T65 which has a germline BRCA1 mutation: a complex rearrangement on chromosome 11 (Supplementary Figure 5a) and a rearrangement involving a translocation between chromosomes 1 and 2 (Supplementary Figure 5b). The mechanisms that give rise to such complex variants are still uncertain, but our observations suggest that these may play a role in the evolution of BRCA-mutated tumors. Further studies are required to ascertain whether BRCA-mutated tumors differ from sporadic breast tumors in the number and type of complex structural variants, as has been characterized for simple structural variants.

## Discussion

Tumors carrying mutations in the BRCA1 and BRCA2 genes, particularly in BRCA1, have more point mutations than sporadic breast tumors, which is not explained by their larger genome size owing to copy number alterations. Mutational signature analysis gives us insight into the number and type of mutational processes acting upon tumor genomes. Here we have used a newly developed method, SparseSignatures, to identify mutational signatures in a previously generated dataset^11^ of 560 breast cancer whole genomes.

SparseSignatures fits a fixed ‘background’ signature representing DNA replication errors, and then discovers additional signatures representing cancer-specific mutational processes. In this dataset, we discover 8 mutational signatures in addition to the background. It is notable that despite finding fewer signatures, our solution still provides a better fit to the data (MSE = 364.345) than the previous solution^11^ with 12 signatures (MSE = 1118.703). Along with providing a better fit to the data, our discovered signatures are also sparser, more clearly differentiated from each other, and lack background noise (Supplementary Table 9).

The previous study on the same dataset^11^ proposed two dense, flat signatures (COSMIC Signatures 3 and 8) to be associated with BRCA1/2 mutated tumors, with a stronger association for signature 3. We do not find either of these signatures in our analysis. While our Signature 8 bears some similarities to the previous ‘Signature 3’, it is considerably sparser and shows stronger nucleotide preferences, which may be due to our explicit separation of the background signature, thus preventing its being confounded with other signatures. We also do not find a signature similar to the previous ‘Signature 30’.

Both this previous study and ours attempted to discover mutational signatures in a dataset of only breast cancers, with the aim of increasing sensitivity to rare signatures which may be present in only a fraction of breast tumors. Another strategy is to discover signatures in a large pan-cancer dataset; this has the advantage of greater accuracy in deciphering signatures if they are present across cancers. Accordingly, we also used a set of 10 signatures that we obtained using SparseSignatures on a pan-cancer dataset of 2827 tumors^21^, and fitted them to the dataset of 560 breast tumors (Supplementary Table 10).

There are two major hypotheses that explain the high risk of cancer in BRCA1/2 mutation carriers; first, homologous repair deficiency leading to elevated and distinct structural variation, and second, cellular dedifferentiation leading to rapid cell division and accumulation of mutations through normal mutagenic processes. If the increased number of mutations in BRCA1/2-mutated tumors was a function of more cell divisions, we would expect this to be explained by higher exposure to the background signature. In fact, we do see higher exposure to the background signature in these tumors, which indicates that they have passed through more cell divisions. However, we also see more mutations attributed to other mutagenic processes, particularly Signature 5 (APOBEC dysregulation leading to C>G mutations) and Signature 8, whose etiology is unknown. This indicates that more cell division may not be the only factor contributing to the higher mutational burden of BRCA1/2-mutated tumors, and that other mutagenic processes are also elevated in these tumors.

Although BRCA1/2-mutated tumors have a higher exposure to the background signature, they do not have a higher exposure to Signature 1, which represents deamination of methylated cytosines at CpG sites. Under conditions of constant DNA methylation, we would expect the exposure values for these two signatures to be proportional to each other. The disproportionately low contribution of Signature 1 to BRCA1/2-mutated tumors suggests a global reduction in methylation levels, which is confirmed by an analysis of TCGA data for BRCA1-mutated tumors. If true, the reduced methylation could cause dysregulation of gene expression and altered binding of gene regulatory proteins. An altered methylation state is also indicative of tumor dedifferentiation, and combined with the indication that these tumors have undergone more cell divisions, further supports the cellular dedifferentiation hypothesis.

It is notable that BRCA1/2-mutated mutated tumors do not appear to possess any unique mutational signatures, suggesting an absence of unique point mutational processes that arise from the BRCA gene mutations. Even Signature 8, which is elevated in BRCA1/2-mutated tumors, also has a high contribution to triple-negative tumors in general. Instead, BRCA1/2-mutated tumors are distinct from all classes of sporadic tumors, including triple-negative tumors, in terms of structural variation. BRCA1/2-mutated tumors have a large number of indels and structural variants, and also present a clearly distinct profile of structural variants. Our analysis confirms a previous finding^26^ that both BRCA1 and BRCA2-mutated tumors have an elevated rate of deletions while BRCA1-mutated tumors specifically have a high rate of tandem duplications. Another previous analysis^11^ sought to identify signatures of structural variation using the same statistical methods as for point mutations. This analysis identified two signatures: ‘rearrangement signature 3’ consisting of tandem duplications 1-100 kb, associated with BRCA1 mutations, and ‘rearrangement signature 5’ consisting of deletions 1-100 kb, associated with both BRCA1 and BRCA2 mutations. Our results are consistent with both of these findings. We also find here that BRCA1-mutated tumors are associated with a higher number of interchromosomal translocations compared to both BRCA2-mutated and sporadic tumors, which to our knowledge has not been shown before.

The functional relevance of structural variants in BRCA1/2 mutated tumors is shown by their enrichment in protein-coding genes, particularly genes with high expression in breast tissue. We identified 11 genes that are enriched for indels and structural variant breakpoints in BRCA1/2-mutated tumors. 4 of these genes (PTEN, RB1, TP53 and RUNX1) are known tumor suppressors and have also been identified as potential point mutation driver genes^23^, showing that in BRCA1/2-mutated tumors, structural variants often inactivate the same drivers that are normally damaged by point mutations in sporadic tumors. Moreover, PTEN, RB1, and TP53 are associated with poor prognosis when inactivated in breast cancer^32,33,34^ and are candidates for targeted therapy^35,36,37^. This suggests that the prevalence of structural mutations in these genes may contribute to the aggressive nature and poor prognosis of BRCA-mutated tumors, and highlights the importance of incorporating structural variant detection into clinical genetic testing^38^. The remaining 6 genes are not known to be enriched for point mutations in breast cancer, and may therefore represent specific indel/structural variant driver genes; of these, ETV6 is known to act as a tumor suppressor in leukemias^39^.

The presence of structural variants disrupting tumor suppressor genes supports the hypothesis that BRCA1/2 tumors may arise due to DNA repair deficiency leading to structural variation. Thus, our study supports both hypotheses and suggests that both of these processes contribute to development of cancer in BRCA mutation carriers.

## Conclusions

Overall, our study suggests that BRCA1/2-mutated tumors are comparatively more aggressive than sporadic breast cancers because loss of the BRCA pathway(s) causes a perfect storm of mutagenic processes and gene dysregulation: less DNA methylation is consistent with the propensity to deregulate and dedifferentiate, and the resulting larger numbers of cell divisions cause a greater point mutational burden; other point-mutagenic processes that may be linked to the tissue of origin and occur in sporadic breast tumors are also active (e.g., APOBEC dysregulation); and crucially, loss of double-strand break repair elevates structural variation rates such that there is a greater chance that driver genes that are hard to functionally affect with point mutations are disrupted at a higher rate than in sporadic tumors.

Given the relatively low number of observed tumours carrying mutations in BRCA1/2 genes, some of the statistical analyses performed through the paper may benefit from future assessments on larger cohorts. As an example, we do not observe significant differences in CpG methylation between BRCA2-mutated tumours vs sporadic ones, but, at the same time, power analysis reveals reduced statistical power for this comparison, and we may be led to different conclusions with larger datasets. The same limitations affect our analysis of structural variants with 10x data. Structural variants may play an important role in characterizing BRCA1/2-mutated tumours and we advocate that future efforts in this direction may shed light on the unique features of these tumours.

## Methods

### Preprocessing data for mutational signature extraction

Point mutations occurring in a genome can be divided into 96 categories based on the base being mutated, the base it is mutated into and its two flanking bases. We therefore represent the dataset of 560 patients from Nik-Zainal et al.^11^ as a mutation count matrix M of size 560 × 96, where element *M*_*i,j*_, is the number of mutations belonging to category *j* in patient *i*. As discussed in SparseSignatures^21^, we removed 5 patients with less than 1000 total mutations, giving a final matrix M of size 555 × 96.

### Modeling mutational signatures

A mutational signature can be represented by a vector *s* of length 96; *s* = [*s*_1_… *s*_96_] where each element *s*_*j*_ represents the probability that this mutagenic process generates a mutation of category *j*. Since these are probabilities, they sum to 1.

Alexandrov et al.^18^ proposed to represent the mutation count matrix *M* as follows:

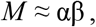

where α_*n* × *K*_ is the exposure matrix (giving the number of mutations contributed by each signature to each patient). α_*ij*_ is the exposure for the *j*^*th*^ signature in the *i*^*th*^ patient. β_*K* × *J*_ is the signature matrix, where each row represents a signature. β_*ij*_ is the proportion of mutations in the *i*^*th*^ signature that fall into the *j*^*th*^ category.

SparseSignatures^21^ incorporates a null model based on mutation rates in the germline. This is the pattern of mutations that would be expected in the course of normal cell division, and is denoted by a vector β_0_ of length *J*, leading to the following representation:

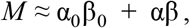

where α_0_ is a vector of exposures, representing the number of mutations contributed by the null (‘background’) signature to each patient.

SparseSignatures also includes two other conceptual improvements: (1) a sparsity constraint based on the LASSO on the matrix β in order to reduce noise and enhance sparsity and separation of the discovered signatures; and (2) a bi-cross-validation approach to choose the number of signatures and avoid overfitting. For details we refer to the paper describing SparseSignatures^21^.

### Implementation of SparseSignatures

In our analysis, we repeated the bi-cross-validation procedure 300 times and we considered values of *K* ranging from 3 to 10 and λ ranging from 0.05 to 0.15. In cross-validation, the configuration with 8 signatures in addition to the background, and λ=0.15, gave the lowest mean squared error on held-out data points. We used the Bioconductor implementation of SparseSignatures (version 1.0.2) in R version 3.3.3.

### Short Read Sequencing

Total genomic DNA was extracted from 14 BRCA+ tumor samples from 13 patients (DNA was extracted from both breasts for one patient) using AllPrep DNA/RNA Mini Kit (Qiagen, Cat. No 80204). The matched control DNA was also isolated from blood of the same patients using Gentra Puregene Blood Kit (Qiagen, Cat. No 158467). To generate short-fragment DNA libraries, 1 ug of total genomic DNA for each sample was sheared to 350 bp. The PCR-free libraries were then constructed from the sheared DNA using Illumina’s TruSeq DNA PCR-Free Sample Preparation Kit. Each library was sequenced with one lane of 2×150bp Illumina HiSeqX sequencing run to 40x genomic coverage.

### 10X Genomics Sequencing

We selected 6 BRCA1/2-mutated tumor samples and 12 sporadic triple-negative tumor samples for 10X genomic sequencing. The long genomic DNA was isolated from 5-10mg tumor core using Gentra Puregene Tissue Kit following the manufacturer’s instructions (Qiagen, Cat. No 158667). Briefly, the small tumor tissue was ground in liquid nitrogen, lysed in Cell Lysis Solution and Proteinase K, and RNA was digested with RNase A. Protein was pelleted and removed by the addition of Protein Precipitation Solution followed by centrifugation. Genomic DNA was precipitated with isopropanol and resuspended in buffer EB. 1.2ng DNA molecules of long fragment were partitioned and barcoded using 10X Genomics Chromium. Each partition had a unique barcode. The barcoded DNA fragments were produced in parallel through emulsion isothermal amplification such that all fragments generated within a partition shared the same barcode. The resulting DNA fragments (Post GEM DNA) from all partitions of the same sample were pooled and recovered. Libraries were constructed following the manufacturer’s protocol through End Repairing, A-tailing, Adaptor Ligation and PCR Amplification. Each library was then sequenced on one lane with a paired-end 150bp run using the Illumina HiSeqX platform to obtain 30x genomic coverage.

### Sequencing data analysis

BWA^40^ v0.7.12 was used to align short-read sequencing data to the human genome. The Long Ranger v1.3 software was used to align 10X genomics data.

### Structural variant calling

BAM files were generated as described above, and for the publicly available data, we downloaded BAM files from the ICGC data portal (https://dcc.icgc.org). We ran SvABA on all BAM files using the default parameters^18^.

Variants with length >=50 bp, as well as interchromosomal translocations, were defined as Structural Variants (SVs) while smaller variants were defined as indels. High-confidence SVs and indels were obtained by selecting variants with (1) both breakpoints in chromosomes 1-22 or X (2) 0 supporting reads in the matched normal sample (3) >=10 supporting reads in the tumor sample, at least 2 of which are split reads in the case of SVs (4) QUAL >= 30 and MAPQ of supporting reads >= 30 (5) neither breakpoint in a gap region (6) Both junctions assembled. We also removed variants that were found in more than one tumor sample or unmatched normal sample, as well as variants found in DGV^41^.

For structural variant detection from 10X genomics prepared samples we used GROC-SVs^30^ with default settings.

### Copy number calling

We used Control-FREEC^27^ to call genome-wide copy number for the samples in our cohort. We used the default parameters for the tool.

### Statistical tests

Numbers of point mutations and structural variants in various groups of samples were compared using the Wilcoxon test. Enrichment of breakpoints in functional regions of the genome was tested using a poisson test, with the null model being that breakpoints are distributed uniformly across the genome (excluding gap regions). To calculate enrichment of indels/SV breakpoints in 10-Mb genomic bins, all the indels and SVs discovered in 46 BRCA-deficient samples were combined and the density of breakpoints across the genome (excluding gap regions) was calculated as 7.36 × 10^−6^/bp. The genome was divided into 574 10-Mb bins overlapping by 5 Mb each. Bins with >25% overlap with gap regions were removed, leaving 516 bins. For each bin, a p-value for enrichment of breakpoints was calculated using a poisson test and bins with a Bonferroni-corrected p-value less than 0.05 were selected. The same procedure was carried out using gene bodies instead of genomic bins to identify genes enriched for breakpoints.

### External data

DNA methylation data, along with clinical data for breast tumor and normal samples, were obtained from The Cancer Genome Atlas. Methylation was measured using the Infinium Human Methylation 450K BeadChip Array from Illumina. The extent of cytosine methylation was represented by a beta value ranging from 0 (fully unmethylated) to 1 (fully methylated), for 482421 CpG and 3081 CpA sites across the genome. Somatic BRCA1 and BRCA2 mutation calls for the same samples were downloaded from cBioPortal (www.cbioportal.org). Germline BRCA1 and BRCA2 mutation calls were obtained with permission from TCGA.

The hg19 human genome was used for all analyses. Positions of genes, exons, open chromatin regions and gap regions were obtained from the UCSC genome browser. Positions of TAD boundaries in MCF-10a cell lines were obtained from Barutcu et al.^42^. Lists of oncogenes and tumor suppressor genes were obtained from the Cancer Gene Census (https://cancer.sanger.ac.uk/census).

Data on replication timing was obtained from Chen et al.^43^. Genomic regions were divided into early-replicating, mid-replicating and late-replicating categories such that a third of the genome for which data was provided was included in each category.

Expression levels in normal human breast tissue was obtained from GTex^29^.

## Declarations

### Ethics approval and consent to participate

Genomic DNA from 14 BRCA1/2-mutated tumors and their matched normal samples was sequenced in this study, as well as genomic DNA from 12 sporadic tumor samples. These samples were obtained as detailed in Telli et al.^24^ This protocol was approved by the institutional review board at Stanford University. Informed consent was obtained from all patients.

### Consent for publication

Not Applicable.

### Availability of data and material

The sequencing data generated during the current study will be made available on dbGAP (accession number will be provided).

Sequencing data from Nik-Zainal et al.^11^ are available from ICGC (https://icgc.org). VCF files and methylation data from TCGA were downloaded from the NCI Genomic Data Commons Legacy Archive (https://portal.gdc.cancer.gov/legacy-archive). GTEx expression data are available from the GTEx Portal (https://www.gtexportal.org/home/datasets). The data used here were based on GTEx analysis v7.

### Authors’ contributions

AS, JMF, AL and DR designed the study. AL and DR analyzed and interpreted sequencing data. ZW carried out preparation and sequencing of tumor samples. KL helped with mutational signature discovery. AL, DR, and AS wrote the manuscript with assistance from JMF. All authors read and approved the final manuscript.

## Acknowledgements

We thank Drs. Noah Spies, Alan Ashworth, David Livingston, and Joan Brugge for discussions. The results published here are based in part upon data generated by the TCGA Research Network (http://cancergenome.nih.gov/) and the Genotype-Tissue Expression (GTEx) Project. The GTEx project was supported by the Common Fund of the Office of the Director of the National Institutes of Health, and by NCI, NHGRI, NHLBI, NIDA, NIMH, and NINDS.

## Supplementary Material Legends

**Supplementary Figure 1.** Boxplots showing the ratio between exposures to Signature 6 (DNA CpG methylation) and the background signature, in BRCA1/2-mutated tumors and various classes if sporadic breast tumors.

**Supplementary Figure 2.** Boxplots showing the average beta-value (extent of DNA cytosine methylation) across 3081 CpA sites, in BRCA1/2-mutated and sporadic breast tumors, based on data from TCGA.

**Supplementary Figure 3.** Boxplots showing the total number of bases in the genome (accounting for copy number changes) in BRCA1-mutated, BRCA2-mutated, and sporadic breast tumors, for the dataset of 81 selected breast tumors.

**Supplementary Figure 4.** Boxplots showing the level of expression in normal breast tissue for genes disrupted by a) indels and b) SVs in 46 BRCA1/2-mutated tumors, based on RNA-Seq data from GTEx^27^. Expression for each gene was measured as median TPM level across all breast tissue samples in the GTEx dataset.

**Supplementary Figure 5.** Two complex structural variants discovered by 10X Genomics sequencing and GROC-SVs in the genome of tumor T65 containing a germline BRCA1 mutation. Inferred extent of breakpoint-supporting read clouds (corresponding to input fragments). The x-axes show chromosomal position. Each row is one read cloud (a cluster of identically barcoded linked reads). The long fragments tile across the breakpoints when ordered by their leftmost position in the left panel.

**Supplementary Table 1.** 9 signatures (including the background) discovered by SparseSignatures on the whole genomes of 560 breast tumors.

**Supplementary Table 2.** Cosine similarities between the 9 signatures (including the background) discovered by SparseSignatures on the whole genomes of 560 breast tumors, and the 30 signatures in the COSMIC database (https://cancer.sanger.ac.uk/cosmic/signatures).

**Supplementary Table 3.** Cosine similarities between the 9 signatures (including the background) discovered by SparseSignatures on the whole genomes of 560 breast tumors.

**Supplementary Table 4.** Exposure values (number of mutations attributed to each signature) for each of the 9 signatures, in each of the 555 breast tumors in which signatures were discovered.

**Supplementary Table 5.** Details of the 771 samples from TCGA used for DNA methylation analysis.

**Supplementary Table 6.** Details of the 81 samples used for structural variant analysis with SvABA.

**Supplementary Table 7.** Details of 10-Mb genomic bins with significant enrichment of structural variant breakpoints, in the combined genomes of 46 BRCA1/2 mutated tumors.

**Supplementary Table 8.** Validation of interchromosomal translocations in BRCA1-mutated tumors using 10X genomics.

**Supplementary Table 9.** Comparison of the 9 signatures (including background) obtained by SparseSignatures with those obtained in a previous study^11^.

**Supplementary Table 10.** Exposure values (number of mutations attributed to each signature) for each of the 10 signatures discovered by applying SparseSignatures to a pan-cancer dataset of 2827 tumors^21^, in each of 560 breast tumors, in the genomes of 555 breast tumors.

