## Supplementary figures and images for "Comprehensive Genomic Characterization of Breast Tumors with BRCA1 and BRCA2 Mutations"

### Supplementary Figure 1

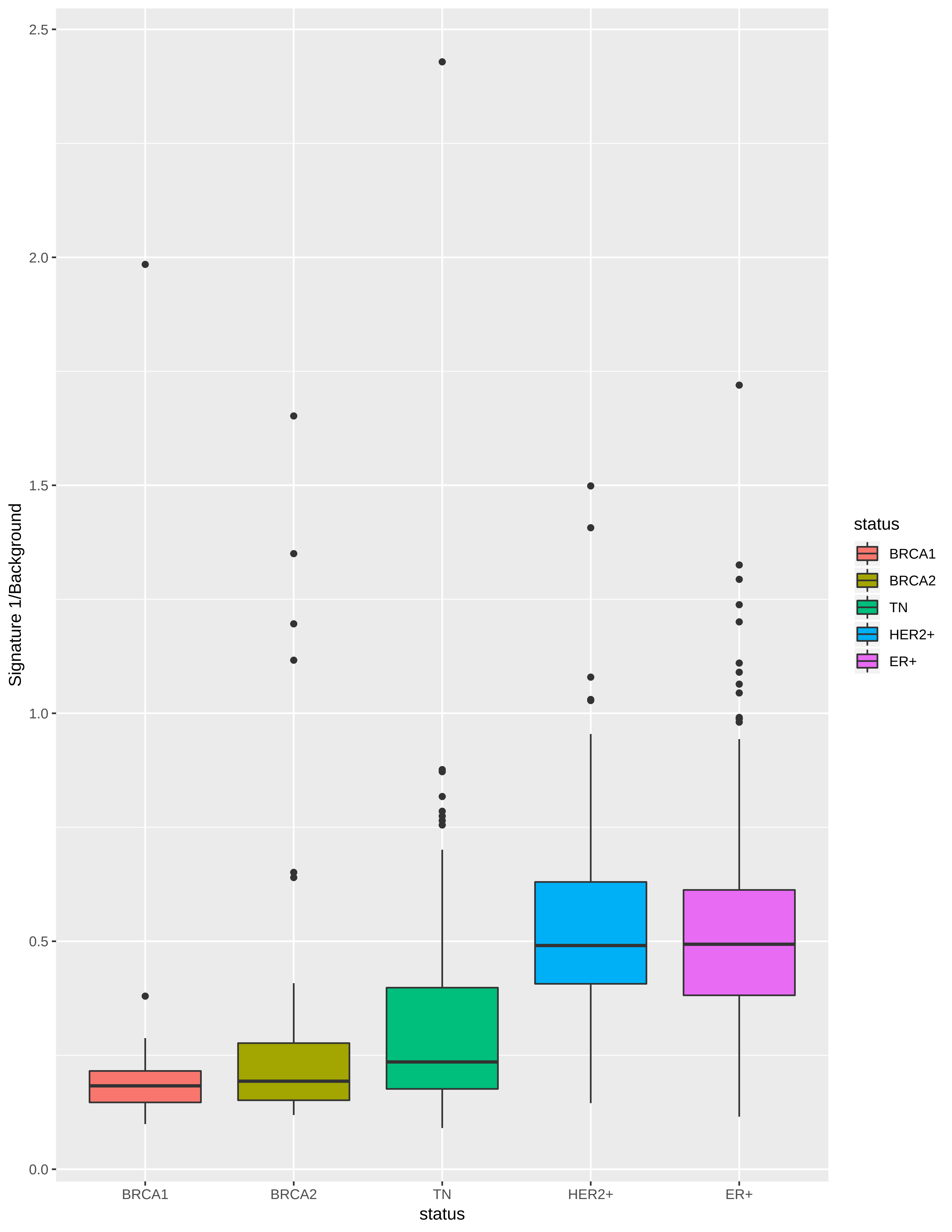

### Supplementary Figure 2

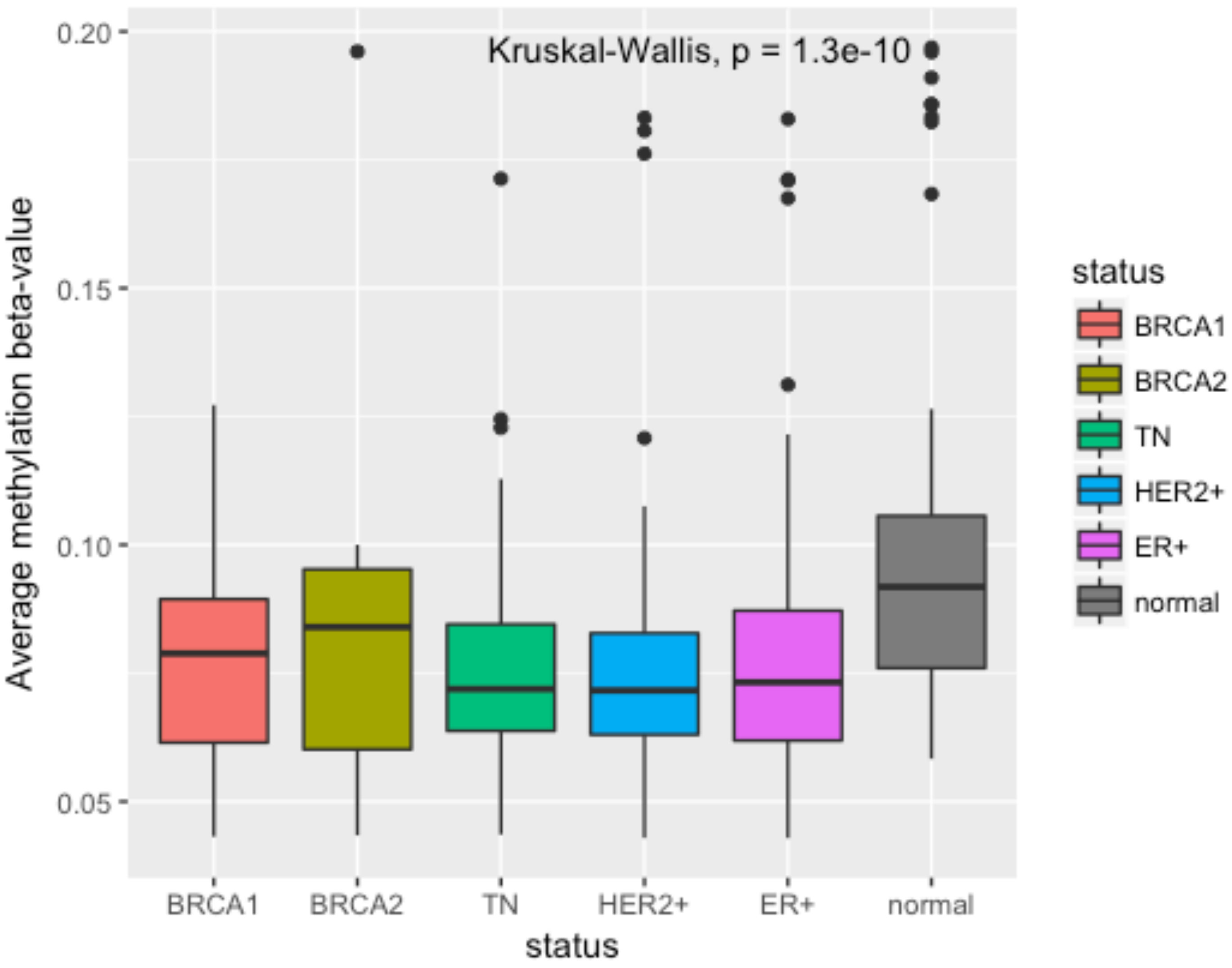

### Supplementary Figure 3

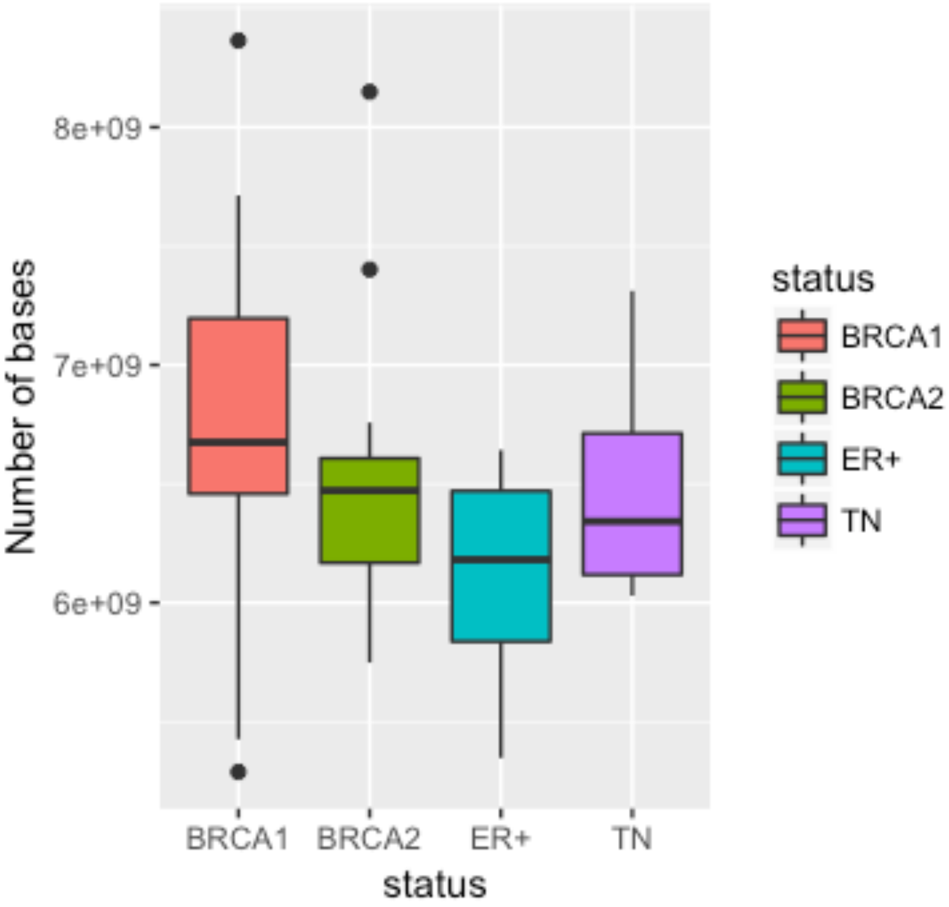

### Supplementary Figure 4

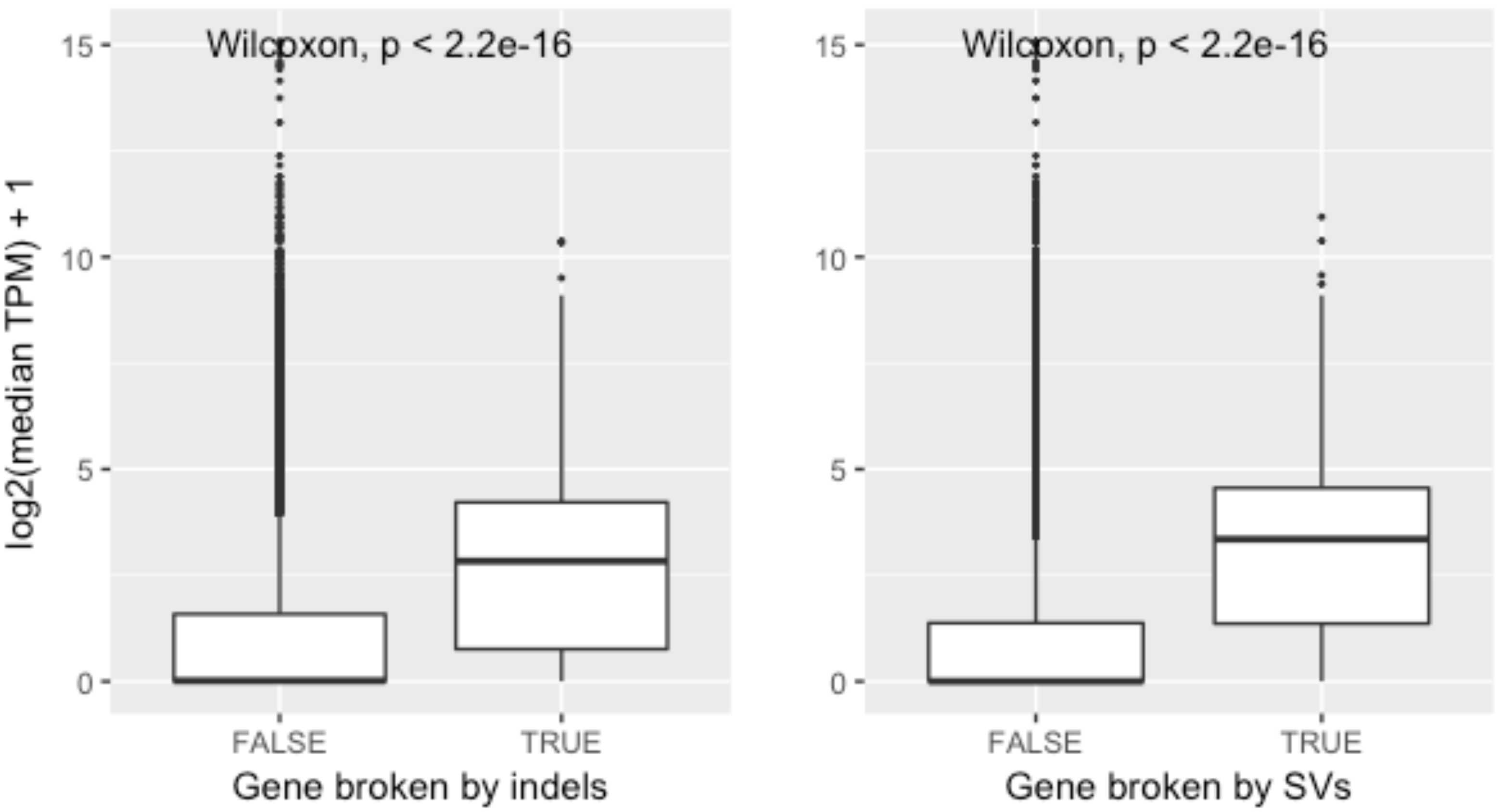

### Supplementary Figure 5

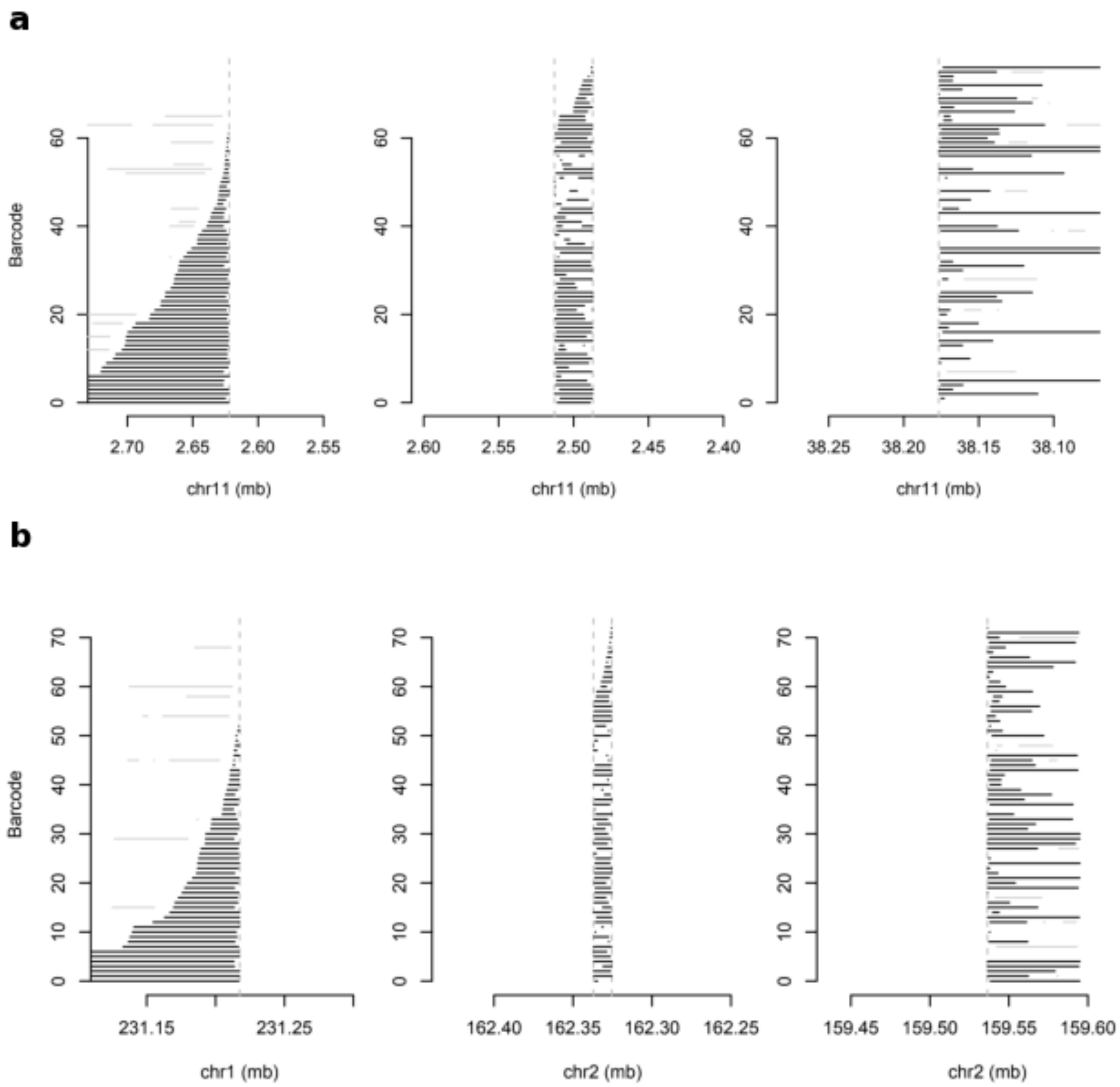
